# Temperature-dependence of metabolic rate in tropical and temperate aquatic insects: support for the Climate Variability Hypothesis in mayflies but not stoneflies

**DOI:** 10.1101/2019.12.25.888578

**Authors:** Alisha A. Shah, H. Arthur Woods, Justin C. Havird, Andrea C. Encalada, Alexander S. Flecker, W. Chris Funk, Juan M. Guayasamin, Boris C. Kondratieff, N. LeRoy Poff, Steven A. Thomas, Kelly R. Zamudio, Cameron K. Ghalambor

## Abstract

A fundamental gap in climate change vulnerability research is an understanding of the relative thermal sensitivity of ectotherms. Aquatic insects are vital to stream ecosystem function and biodiversity but insufficiently studied with respect to their thermal physiology. With global temperatures rising at an unprecedented rate, it is imperative that we know how aquatic insects respond to increasing temperature and whether these responses vary among taxa, latitudes, and elevations. We evaluated the thermal sensitivity of standard metabolic rate in stream-dwelling baetid mayflies and perlid stoneflies across a ~2,000 m elevation gradient in the temperate Rocky Mountains in Colorado, U.S.A., and the tropical Andes in Napo, Ecuador. We used temperature-controlled water baths and microrespirometry to estimate changes in oxygen consumption. Tropical mayflies generally exhibited greater thermal sensitivity in metabolism compared to temperate mayflies; tropical mayfly metabolic rates increased more rapidly with temperature and the insects more frequently exhibited behavioral signs of thermal stress. By contrast, temperate and tropical stoneflies did not clearly differ. Varied responses to temperature among baetid mayflies and perlid stoneflies may reflect differences in evolutionary history or ecological roles as herbivores and predators, respectively. Our results show that there is physiological variation across elevations and species and that low elevation tropical mayflies may be especially imperiled by climate warming. Given such variation among species, broad generalizations about the vulnerability of tropical ectotherms should be made more cautiously.

## Introduction

Metabolism integrates an organism’s existing energy supplies and underlies fitness-related activities like movement, growth, tissue maintenance, and development (Dillon, Wang, & Huey, 2010; Gillooly, Brown, West, Savage, & Charnov, 2001). In ectotherms, metabolic rates are sensitive to body temperature, which can influence not only individual fitness, but also geographic distributions and abundances (Angilletta, 2009; Terblanche & Chown, 2007; Vannote & Sweeney, 1980). A major challenge in global change biology is to understand how temperature alters metabolism and energy budgets and, in turn, the ability of species to cope with climate change (Colwell, Brehm, Cardelús, Gilman, & Longino, 2008; Dillon et al., 2010; Parmesan & Yohe, 2003; Walther et al., 2002). To date, attempts to measure vulnerability to climate change have largely assessed species responses to temperature challenges, such as lethal and critical thermal limits (Deutsch et al., 2008; Pinsky, Eikeset, McCauley, Payne, & Sunday, 2019). Despite their utility, these metrics do not capture the continuous or incremental sub-critical effects of temperature on an organism’s energy budget. Such effects are particularly important in ectotherms, because body temperature and metabolic rate increase with environmental temperature, resulting in higher energy expenditure. This response is likely to trade off against other demands on energy budgets, such as growth, reproduction, and locomotor performance (Kozłowski, 1992; Pörtner et al., 2006; Wieser, 1994). For example, in aquatic ectotherms, increased metabolic demand resulting from higher temperature can outpace the supply of oxygen from the environment causing decreased performance and lowered tolerance to heat stress (Pörtner, Bock, & Mark, 2017). Thus, understanding how metabolic responses to temperature vary across different thermal environments can help us predict geographic variation in response to global warming (Dillon et al., 2010).

In this study, we measured metabolic rate profiles in juveniles (nymphs) of two major groups of stream insects – mayflies and stoneflies – in temperate and tropical streams across a paired elevation gradient. Mayflies and stoneflies spend most of their life cycles as aquatic nymphs, during which they feed, grow, and develop eggs and sperm before emerging to breed as short-lived flying adults (Brittain, 1990). Water temperature is therefore a major driver of the life cycle in these taxa. We aimed to elucidate differences in patterns of species’ thermal sensitivity and thus better evaluate potential vulnerabilities of mayflies and stoneflies to projected climate warming (Chown, Duffy, & Sørensen, 2015).

Much like other types of thermal performance curves (TPCs), oxygen-consumption profiles (used to measure metabolic rate) vary with increasing temperature and display three identifiable regions (Pörtner, 2002; Schulte, 2015): 1) an ascending phase; 2) a peak, which indicates the onset of stress (Huey & Stevenson, 1979; Pörtner & Knust, 2007); and 3) a descending phase of metabolic depression (Guppy & Withers, 1999). When coupled with other measures of performance and critical thermal limits, changes in oxygen consumption provide a robust picture of how temperature shapes the ecology of an organism (Angilletta, 2009; Pörtner et al., 2017) (Fig. 1).

**Figure 1.**
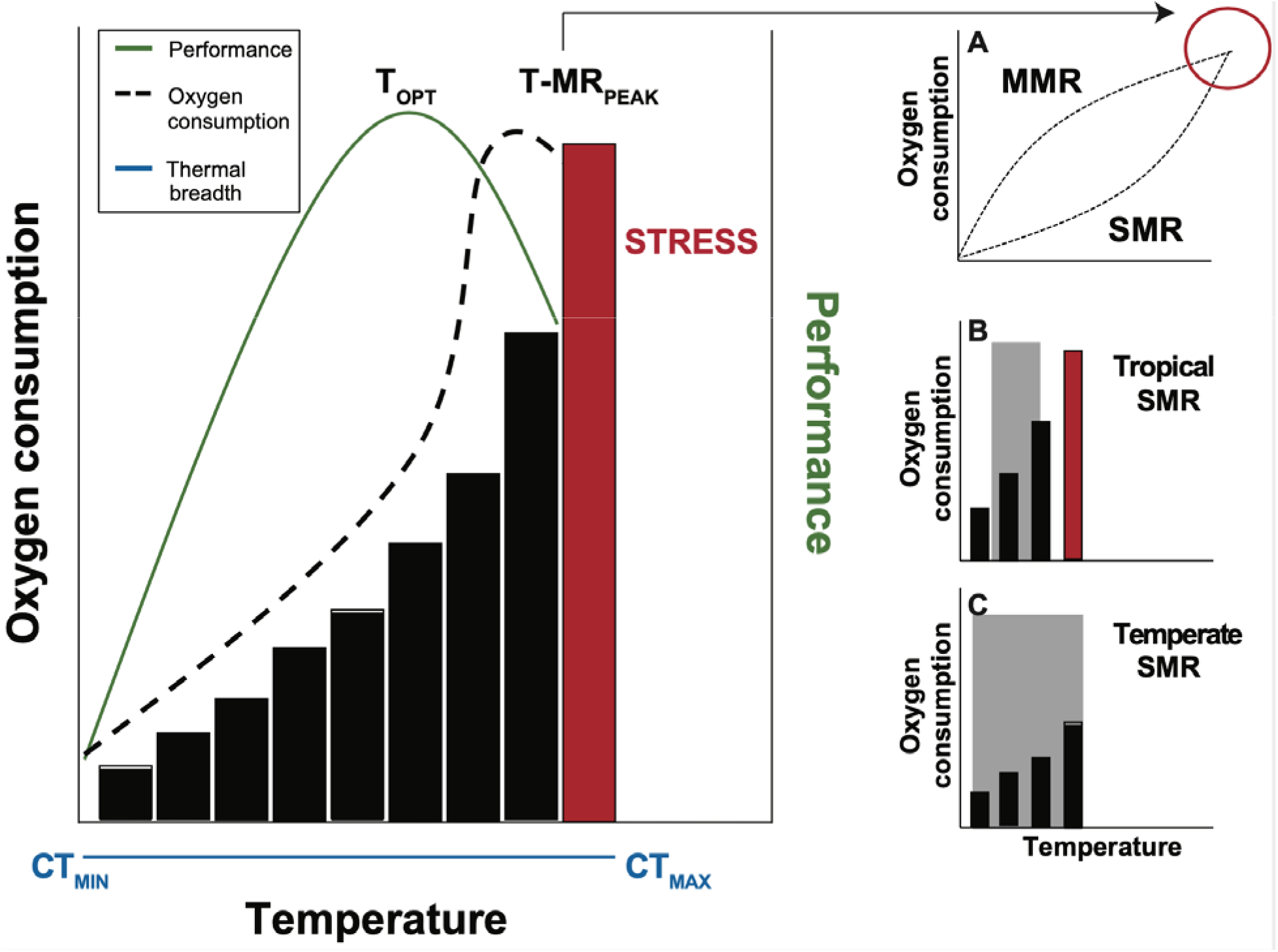
Expectations for change in standard metabolic rate (oxygen consumption) and performance in response to temperature in an aquatic ectotherm. The thermal performance curve (TPC, solid green line, left panel) refers to any functional performance trait (e.g., swimming performance), where higher performance is better. T_OPT_, where performance is highest, reflects the animal’s preferred temperature. The oxygen consumption curve (dashed line) is a special case of TPC. The peak in this case (T-MR_PEAK_) could represent the temperature at which standard metabolic rate (SMR) is so high that it essentially equals the maximum metabolic rate (MMR) that the ectotherm can achieve (circle, A). T-MR_PEAK_ may therefore reflect stress (red) and aligns with a decline in the functional performance trait. The general location of behaviorally determined critical thermal limits (CT_MIN_ and CT_MAX_) are also shown (blue line, left panel). In this study we focus only on SMR. We predict that SMR will rise steeply in tropical species (B) and be stressful when temperatures exceed the typical narrow range of temperatures they experience (gray rectangle). In comparison, temperate species should exhibit a more gradual rise in SMR across the same temperatures (C), reflecting the broad range of environmental temperatures they experience (grey rectangle).

Although TPCs for traits like locomotion and metabolic rate are expected to have similar shapes, a metabolic rate curve is a special type of TPC (Pörtner, 2002; Schulte, Healy, & Fangue, 2011), with different implications for organismal fitness (Kellermann et al., 2019). The performance traits used in many TPCs—running speed, feeding rates, or growth—are often interpreted as components of fitness, such that *more is better*. For these traits, peaks indicate the location of thermal optima (T_OPT_, Fig. 1) and suggest that performance is optimized at that temperature (Angilletta, Niewiarowski, & Navas, 2002; Huey & Kingsolver, 1989; Huey & Stevenson, 1979) or nearby (Martin & Huey, 2008). For metabolism, however, more is not necessarily better, and peak rates of standard oxygen consumption represent elevated minimum costs, typically occurring at warmer temperatures than peak performance traits (Pörtner, 2001). For example, aerobic scope is maximized where the difference between standard metabolic rate (SMR) and maximum metabolic rate (MMR) is greatest (Fig. 1A; often the location of peak performance T_OPT_) because maintenance costs are minimized relative to the peak demand (Clark, Sandblom, & Jutfelt, 2013). At higher temperatures, when SMR meets MMR, a metabolic depression often follows as organismal functions decline (Fig. 1; also see Pörtner, 2002). Presumably, these temperatures lie above the maximum temperatures typically experienced by the organism and are likely to be stressful. Thus, the metabolic peak should lie to the right of T_OPT_ (i.e., at a higher temperature than T_OPT_), align with decreasing performance on the TPC, and should be interpreted as the onset of stress (Fig. 1; Pörtner, 2001). We refer to this peak as T-MR_PEAK_.

The specific shapes of metabolic rate curves found for different species, populations, and thermal experiences (e.g., acclimation; DeLong et al., 2018; Havird et al., 2020) could reflect either the acute effects of temperature on biological rates (Dillon et al., 2010; Payne & Smith, 2017) or the evolutionary response to past climatic conditions (Dell, Pawar, & Savage, 2011; Deutsch et al., 2008). If the shape of the metabolic rate TPC simply reflects the thermodynamics of biological reactions, we would predict that organisms of equivalent size experiencing a similar range of temperatures should exhibit similar metabolic rate curves regardless of their evolutionary history (Payne & Smith, 2017).

Alternatively, the Climate Variability Hypothesis posits that if stable climatic regimes, such as those found in the tropics, expose organisms to only a narrow range of temperatures and favor the evolution of thermal specialists (*sensu* Huey & Kingsolver, 1989), then we predict that those organisms should have lower SMR within the range of temperatures they experience to maximize the allocation of energy to growth, reproduction, and aerobic performance (Fig. 1B). These thermal specialists should also be more sensitive to temperature, i.e., have metabolic rates that change rapidly in response to temperature outside their normal range. Climates characterized by large seasonal temperature variation, such as those found at temperate latitudes, should select for thermal generalists capable of functioning effectively over a wide range of temperatures (Angilletta et al., 2002; Gilchrist, 1995; Huey & Kingsolver, 1993). Thermal generalists are predicted to have relatively low SMRs across a wider range of temperatures (Fig. 1C) and be less sensitive to increasing temperature to keep their minimum energy demands low. The generality of these contrasting hypotheses remains unresolved because few comparative studies have measured the thermal sensitivity of metabolic rates in species from temperate and tropical regions under similar experimental conditions.

To test predictions generated by these alternative hypotheses, we compared standard metabolic rates of temperate and tropical mayflies and stoneflies across different elevations. Because stream temperatures at a given elevation were similar at the time of the experiments, we predicted that the metabolic rate curves should look similar between temperate and tropical species if they were strictly determined by thermodynamic effects on metabolic processes (Payne & Smith, 2017). Alternatively, if metabolic rates reflect adaptation to climate, we predicted greater sensitivity to temperature in the tropical species (Fig. 1B) compared to temperate ones (Fig. 1C; Angilletta et al., 2002; Huey & Kingsolver, 1993).

## Methods

### Site and species descriptions

Temperate sites were located in the Cache La Poudre river drainage of the southern Rocky Mountains in Colorado, USA, and tropical sites in the Papallacta-Quijos river drainage of the Andes Mountains in Napo Province, Ecuador. We chose small, relatively pristine streams where the mayflies and stoneflies used in this study are typically found. Sites at both latitudes were matched by elevation, where high >3,000 m, mid = 2,500 – 3,000 m, and low = 1,000 – 2,500 m. Streams within each elevation pair were approximately the same temperature at the time of collection (Table 1). Thus, we were able to make latitudinal comparisons of insect metabolic rate for each elevation pair under the same thermal acclimation conditions.

**Table 1.** Average stream temperatures with standard errors and acclimation temperatures for each temperate (Rocky Mountain) and tropical (Andes) stream from which insects were collected.

| Latitude | Site ID | Type | Elevation (m) | Stream temperature (°C) | Acclimation temperature (°C) |
| --- | --- | --- | --- | --- | --- |
| Temperate | COP 2212 | Low | 2212 | 12.5 ± 3.73 | 14.0 |
|  | COP 2590 | Mid | 2590 | 10.8 ± 4.68 | 10.0 |
|  | COP 2798 | Mid | 2798 | 7.6 ± 1.41 | 10.0 |
|  | COP 3166 | High | 3166 | 6.2 ± 3.56 | 6.0 |
| Tropical | ECP 1845 | Low | 1845 | 13.5 ± 0.72 | 14.0 |
|  | ECP 2003 | Low | 2003 | 14.0 ± 0.38 | 14.0 |
|  | ECP 2694 | Mid | 2694 | 10.0 ± 0.79 | 10.0 |
|  | ECP 3683 | High | 3683 | 8.6 ± 0.89 | 6.0 |
|  | ECP 3898 | High | 3898 | 6.6 ± 1.74 | 6.0 |

Ephemeroptera (mayflies) and Plecoptera (stoneflies) do not share genera between tropical and temperate sites. Therefore, we selected representative species from the same family (i.e., mayfly family Baetidae and stonefly family Perlidae) at each location to contrast their metabolic rate. Based on our previous DNA-barcoding work (Gill et al., 2016; Polato et al., 2018; Shah et al., 2017b), we could identify taxa to species using morphological descriptions of individuals from these sites. Ultimately, we chose these taxa because 1) they represent the most closely related phylogenetic temperate and tropical relatives, 2) they are ecologically and morphologically very similar to each other, and 3) they tend to be common in streams around the world. Results from our work could therefore have general implications.

In the Rockies, we collected 313 mayflies from the genus *Baetis* (*Baetis bicaudatus* at high and mid elevations and *Baetis tricaudatus* at low elevations) and 107 stoneflies (*Hesperoperla pacifica* at low and mid-elevations). In the Andes, we collected 245 mayflies from the genus *Andesiops*. Earlier work has shown cryptic diversity in this genus, such that *Andesiops* collected at each elevation were putatively different species (Gill et al., 2016). We collected 76 stoneflies from the genus *Anacroneuria*, which comprises *Anacroneuria rawlinsi* at low elevation and *Anacroneuria guambiana* at mid-elevation. Although baetid mayflies were easily collected across all elevations, perlid stoneflies were much harder to find and were altogether absent at high elevation, so latitudinal comparisons between stoneflies are at mid and low elevations only. We conducted experiments between January and April of 2013 in the Andes and June to August of 2013 in the Rockies.

### Lab acclimation

Field-caught nymphs were brought back to the lab (~1,600 m at both latitudes) and placed in containers of filtered stream water from their native streams without food for 48 h. During this time, high oxygen levels were maintained by bubbling air into the containers, and water was circulated by small pumps, with temperatures approximating the insects’ home stream temperatures (Table 1). Specifically, acclimation temperatures were chosen to closely match the average temperature of the temperate and tropical stream pair at each elevation. Temperate and tropical high elevation insects were acclimated to 6.0 °C, mid-elevation insects to 10.0 °C, and low elevation insects to 14.0 °C (Table 1). Acclimation temperatures did not exactly match average stream temperatures (Table 1) due to the difficultly of maintaining precise temperatures in large water baths with our heating and cooling equipment, especially in the tropics. Notwithstanding these limitations, the difference in acclimation versus average stream temperature was between 0 - 2.6 °C at any given site. We selected a paired elevation design because tropical insects failed to acclimate to temperatures outside average temperatures of their native streams (see also Shah, Funk, & Ghalambor, 2017a). For example, we often detected evidence of stress (e.g., jerking movements, unnatural swimming patterns) and even mortality, especially when tropical mayflies from high and low elevations were warmed for 48 h outside native stream temperature (i.e., 12 °C and 10 °C, respectively). Therefore, the use of a single acclimation temperature across all elevations was not possible. Because acclimation temperatures differed among elevations but were the same for each elevation pair, we restricted our statistical comparisons to the same elevations between the two latitudes.

### Metabolic rate

Oxygen consumption rates (metabolic rates, hereafter) were measured using closed glass micro-respirometry chambers (Unisense A/S, Aarhus, Denmark). After the 48 h acclimation period, we introduced insects randomly to one of nine test temperatures (5, 7.5, 10, 12.5, 15, 17.5, 20, 22.5, or 25 °C) by allowing temperature to rise naturally or by slowly cooling (0.5 °C · min^−1^) to prevent thermal shock. On average, 16 individuals per population for mayflies (min = 3, max = 22) and 6 per population for stoneflies (min = 3, max = 12) were tested at each temperature (Tables S1, S2). Each individual was tested at only one temperature. Once at the test temperature, insects were moved individually into glass chambers filled with stream water that had been treated with UV light to reduce background microbial respiration and immersed in a water bath held at the test temperature (Fig. S1). Mayflies were placed in 4 mL chambers, and stoneflies, which are larger, were placed in 50 mL chambers. Chambers were fitted with a glass-coated magnetic stir bar and a plastic mesh to which the insect could cling (Fig. S1). We acclimated insects to the chamber space for an additional hour before sealing each chamber with a glass stopper to prevent further introduction of atmospheric oxygen. Each stopper had a central capillary hole, allowing us to insert an oxygen microelectrode (500 μm diam.) connected to a picoammeter (Unisense OXY-Meter, Aarhus, Denmark) to measure change in oxygen concentration. We measured background respiration rates by placing a control chamber (without an insect) in each experiment. The microelectrode was first lowered into the control chamber at the start of the experiment. After reading oxygen concentration continuously for 2 mins, it was moved to the first chamber that contained an animal, where data were recorded for another 2 mins (Fig. S1), and so on. Experiments were terminated when oxygen levels reached 80% of the air-saturated values because natural dissolved oxygen levels in our study sites rarely fall below this value. Consequently, we made 3-6 readings for each chamber – fewer readings at the higher temperatures due to the combined effect of lower oxygen concentrations and faster oxygen consumption by insects – which lasted around 128 minutes. Because there were typically 16 chambers in each experiment, the average time interval between readings was 25 mins. While recording oxygen concentration, we also observed insects inside the chambers, noting any visible behavioral stress responses or mortality. Insects were then removed from chambers, euthanized in ethanol, dried, and weighed on a laboratory scale with readability to 0.01mg.

### Statistical Analyses

We first calculated average rate of oxygen consumption for each insect or control (blank) chamber by measuring change in oxygen concentration across all measurements taken in a given chamber (see Supplementary Materials and Fig. S2 for an explanation of this calculation). We then averaged the control values for each temperature such that we calculated a control rate for each combination of test temperature, elevation, and latitude. The mean control rates were subtracted from individual insect consumption rates to correct for background microbial respiration at that temperature. We conducted all statistical analyses on these ‘control-corrected oxygen consumption rates’ (hereafter, oxygen consumption) in R version 3.4.0 (R Core Team 2017). For each taxon, latitudinal contrasts were conducted separately for each elevation category. Thus, for mayflies, we compared temperate and tropical baetids in three statistical comparisons: low, mid, and high elevation. Similarly, for stoneflies, we ran two separate analyses for low and mid elevation.

Because we were interested in the thermal sensitivity of temperate and tropical insect metabolic rates between elevation pairs (i.e., the thermal reaction norm), we first compared metabolic rates during the ascending phase of the TPC by fitting a linear model with oxygen consumption as the response variable for each elevation type (i.e., high, mid, and low) separately. For this analysis, we used only the temperatures 5, 7.5, 10, 12.5, 15 and 17.5 °C because these temperatures were within the ascending phase of the SMR curves for all populations in this study. The model included fixed effects of test temperature and latitude, and their interaction, with body mass as a covariate (Packard & Boardman, 1999). A significant interaction would denote a difference in thermal sensitivity (i.e., slope) between temperate and tropical species.

We conducted a similar analysis for the descending phase using data from the 17.5 and 20 °C treatments as these two temperatures encompassed the descending phase of the curve. At the highest temperatures (22.5 and 25 °C), we observed stress responses (e.g., unnatural movement, frenzied attempts to escape from the chamber and in some cases elevated metabolic rates) in tropical mayflies. We therefore did not include SMR at 22.5 and 25 °C in this analysis but instead calculated separate Q_10_ values for these temperatures (see below).

To compare the maximum SMR at each elevation between temperate and tropical taxa, we examined the latitude x elevation combination individually to find the physiologically relevant peak (i.e., before CT_MAX_ and when the first signs of stress were apparent, if observed). This peak was typically at 17.5 °C in our study because in most cases, across both latitudes, SMR was highest at this temperature even if not always statistically different from 15 °C. We then compared these values using an ANOVA for each elevation pair using latitude as the fixed effect and body mass as a covariate. For all analyses, we plotted the least square means to control for the effect of body mass on standard metabolic rates.

Finally, we calculated Q_10_, a measure of sensitivity to temperature, using the formula

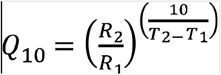

where R_1_ and R_2_ are reaction (oxygen consumption) rates measured at two different temperatures, T_1_ and T_2_, respectively. 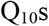 were calculated across cooler, ‘non-stressful’ temperatures (T_1_ = 7.5 and T_2_ = 15 °C) to ensure measurements were within the exponential phase of the metabolic curve and warmer, ‘stressful’ temperatures (T_1_ = 15 and T_2_ = 25 °C) to capture the relative decline in rates.

## Results

### Behavioral observations

Tropical mayflies showed visible signs of stress (jerking movements, loss of righting response) at several test temperatures outside the range typically experienced (e.g., 22.5 and 25 °C; Fig. 2 red bars, Table S1; Fig. S3A); many of these visibly stressed individuals died at the end of the experiment (Table S.1; Fig. S3B). Low elevation tropical mayflies also experienced some mortality at 5 °C, which is much colder than the minimum temperatures they encounter in their home streams (Fig. 2F). Temperate mayflies, by contrast, did not show visible signs of stress at any experimental temperature (Table S2). We also did not detect any overt signs of stress in any of the tropical or temperate stoneflies in this study.

**Figure 2.**
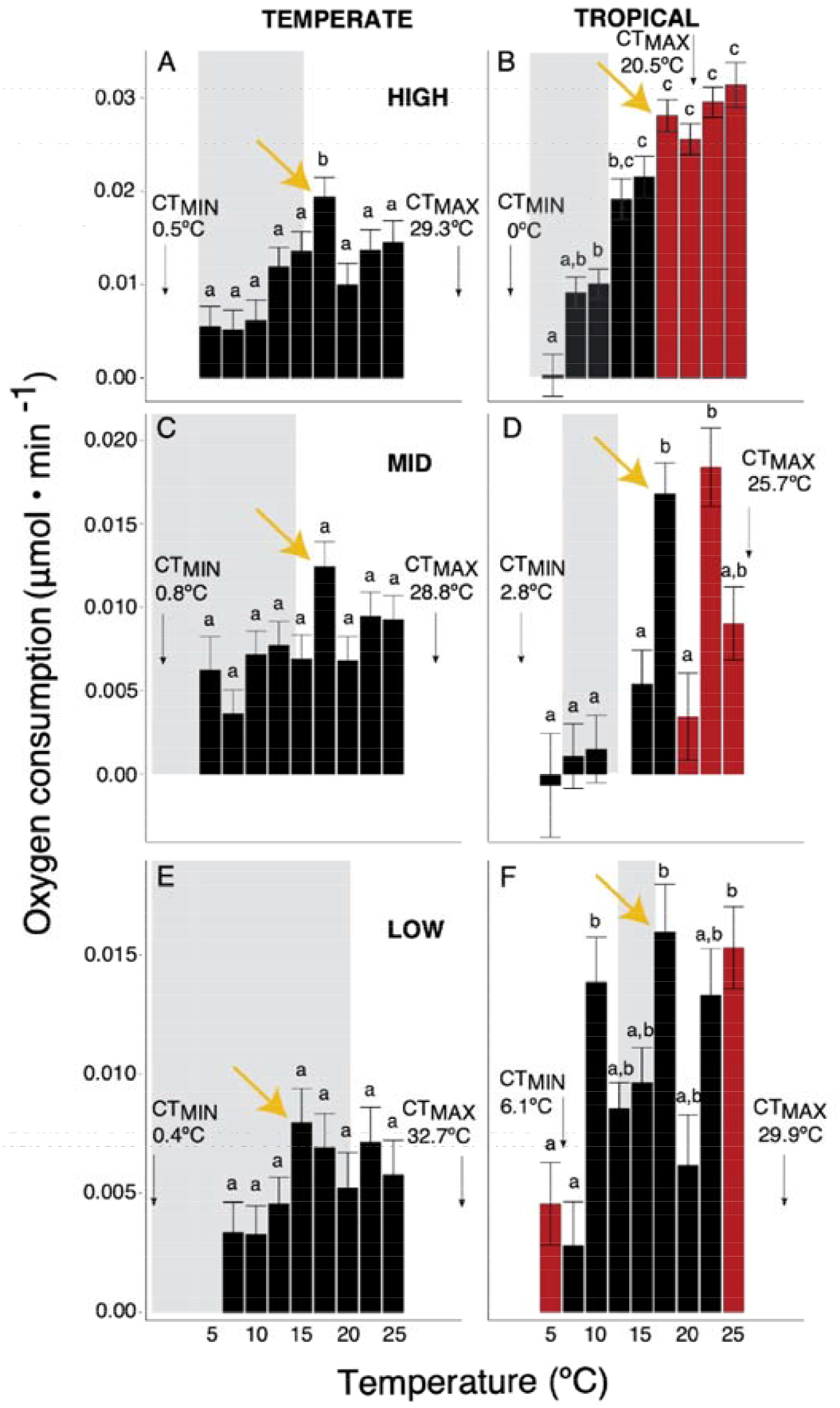
Plots of standard metabolic rate least square means in baetid mayflies from high (A, B), mid (C,D), and low (E,F) elevations in temperate (left panel) and tropical (right panel) streams. Red bars indicate temperatures at which we observed stress. T-MRPEAK is shown with yellow arrows for each population. Gray rectangles represent the annual stream temperature range measured in each stream.

### Thermal sensitivity of standard metabolic rates and reaction norms

SMR profiles differed between temperate and tropical mayflies and stoneflies at many elevations (Figs. 2, 3). In general, metabolic rates of mayflies from the two latitudes were lowest at temperatures approximating those in their native streams. Outside these temperatures, however, tropical mayflies exhibited a greater increase in metabolic rate (Fig. 2). In all mayfly populations across both latitudes, a secondary peak was observed at 22.5 and 25 °C that corresponded with visible stress during experiments. We denoted the initial peak in SMR as T-MR _PEAK_ (shown as yellow arrows on Figs. 2 and 3) or the onset of stress, which occurred at temperatures higher than maximum stream temperature, as expected. Secondary peaks were viewed as an acute stress response because they were followed by death in some cases. As indicated in the Methods, we excluded these stressful test temperatures from the analyses of the ascending and descending phases of metabolic rate profiles and dealt with them separately in an analysis of Q_10_.

**Figure 3.**
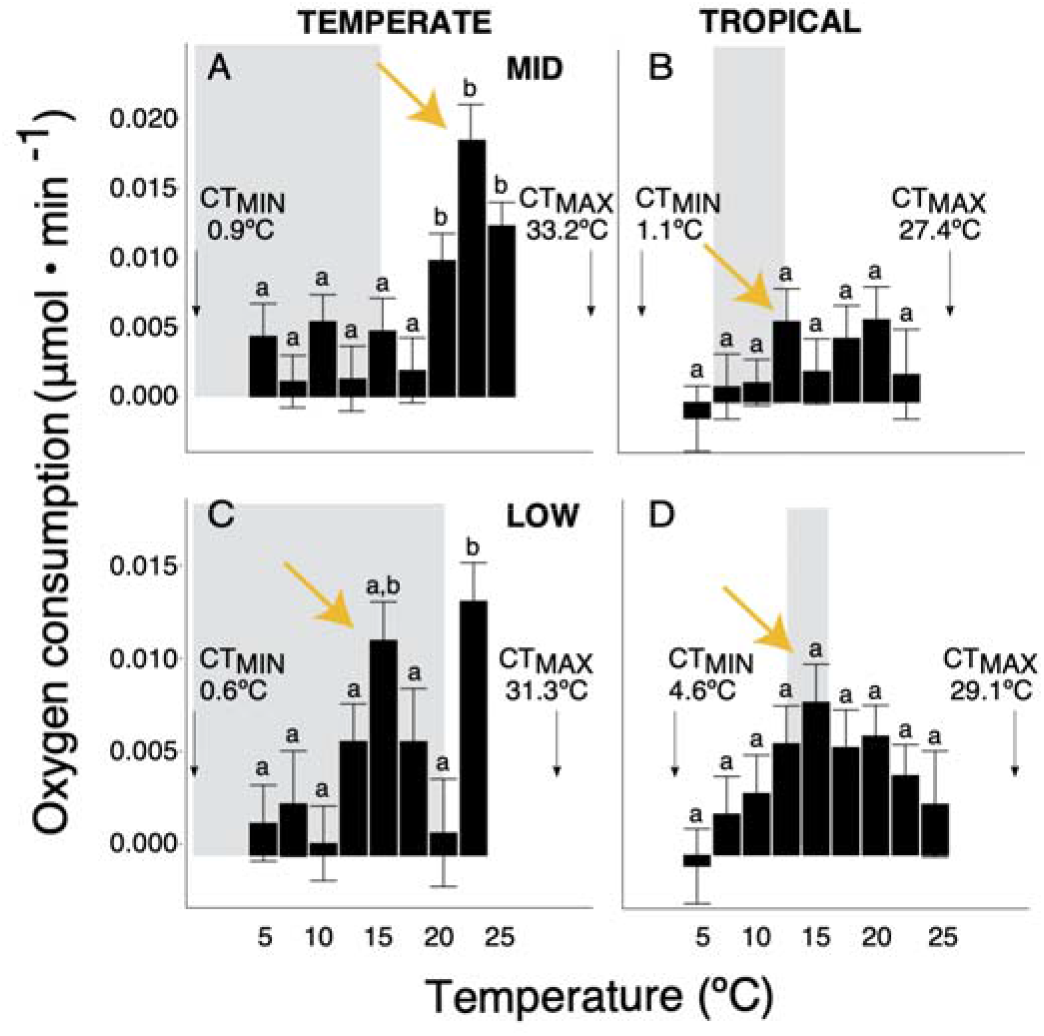
Plots of metabolic rate least square means in perlid stoneflies from mid (A, B) and low (C,D) elevations in temperate (left panel) and tropical (right panel) streams. Stoneflies were not found at high elevation. T-MR_PEAK_ is shown with yellow arrows for each population. Gray rectangles represent th approximate stream temperature range measured in each stream. We did not detect overt stress in stoneflies in this study. CT_MIN_ and CT_MAX_ determined in a previous study are shown.

In the ascending phase, metabolic rates of tropical mayflies were more sensitive to temperature increases than their temperate counterparts at high elevation (*F*_(1,105)_ = 13.79, *p* < 0.001, Fig. 4A) and mid elevation (*F*_(1,105)_ = 11.95, *p* = 0.001, Fig. 4B). At low elevation, despite high mortality and visible signs of stress in tropical but not temperate mayflies, we did not find a significant interaction between tropical and temperate groups (*F*_(1,159)_ = 1.505, *p* = 0.222, Fig. 4C). In stoneflies, ascending phases did not differ between temperate and tropical populations (mid elevation: *F*_(1,79)_ = 2.363, *p* = 0.128; Fig. 5A and low elevation: *F*_(1,61)_ = 0.136, *p* = 0.714; Fig. 5B)

**Figure 4.**
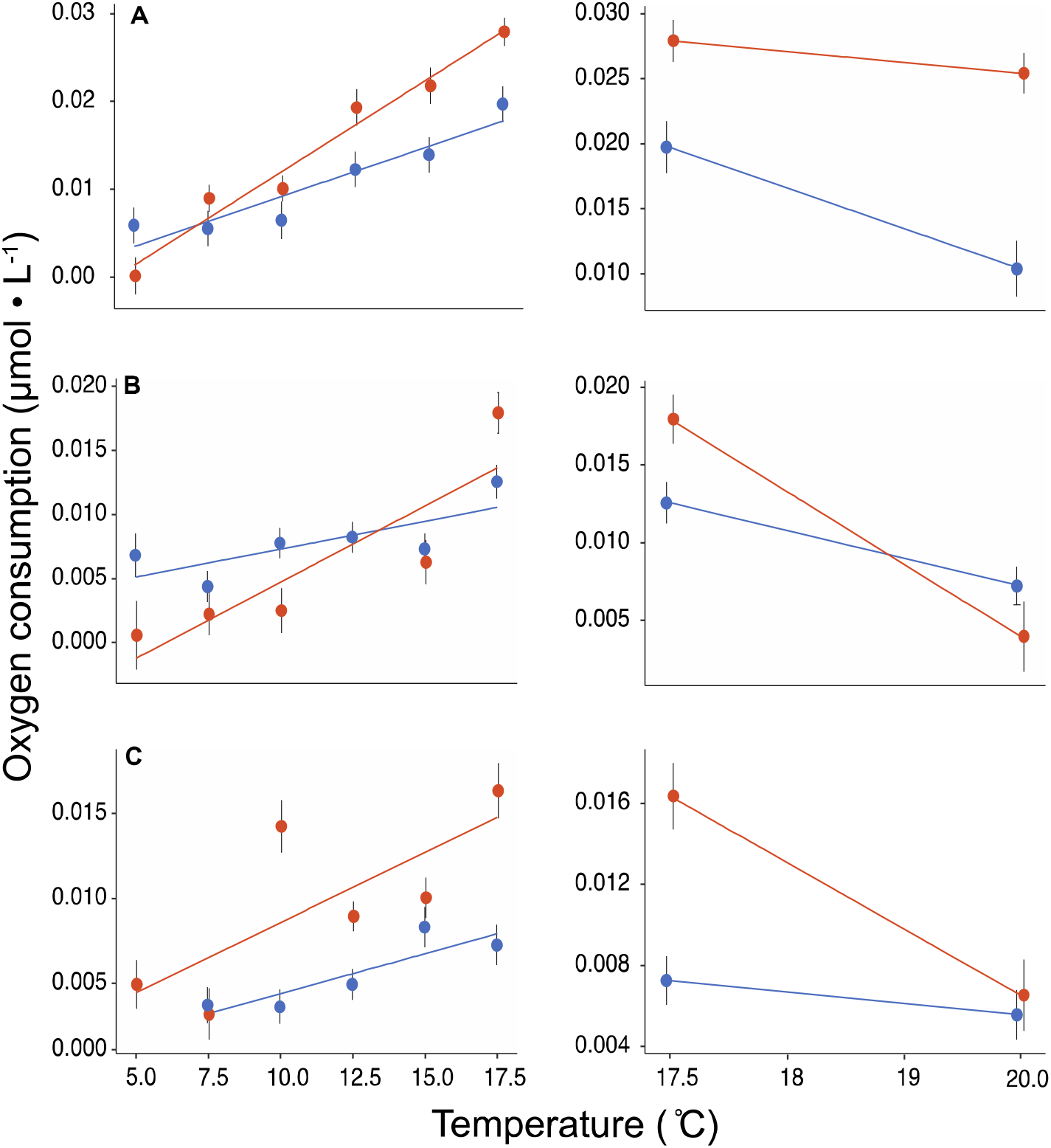
A comparison of the ascending (left panel) and descending (right panel) phases of metabolic profiles in baetid mayflies at high (A), mid (B) and low (C) elevations. Plotted values are the least square means of metabolic rate in mayflies from tropical (orange circles) and temperate (blue circles) streams, after accounting for body mass in the model. Vertical bars are standard errors.

**Figure 5.**
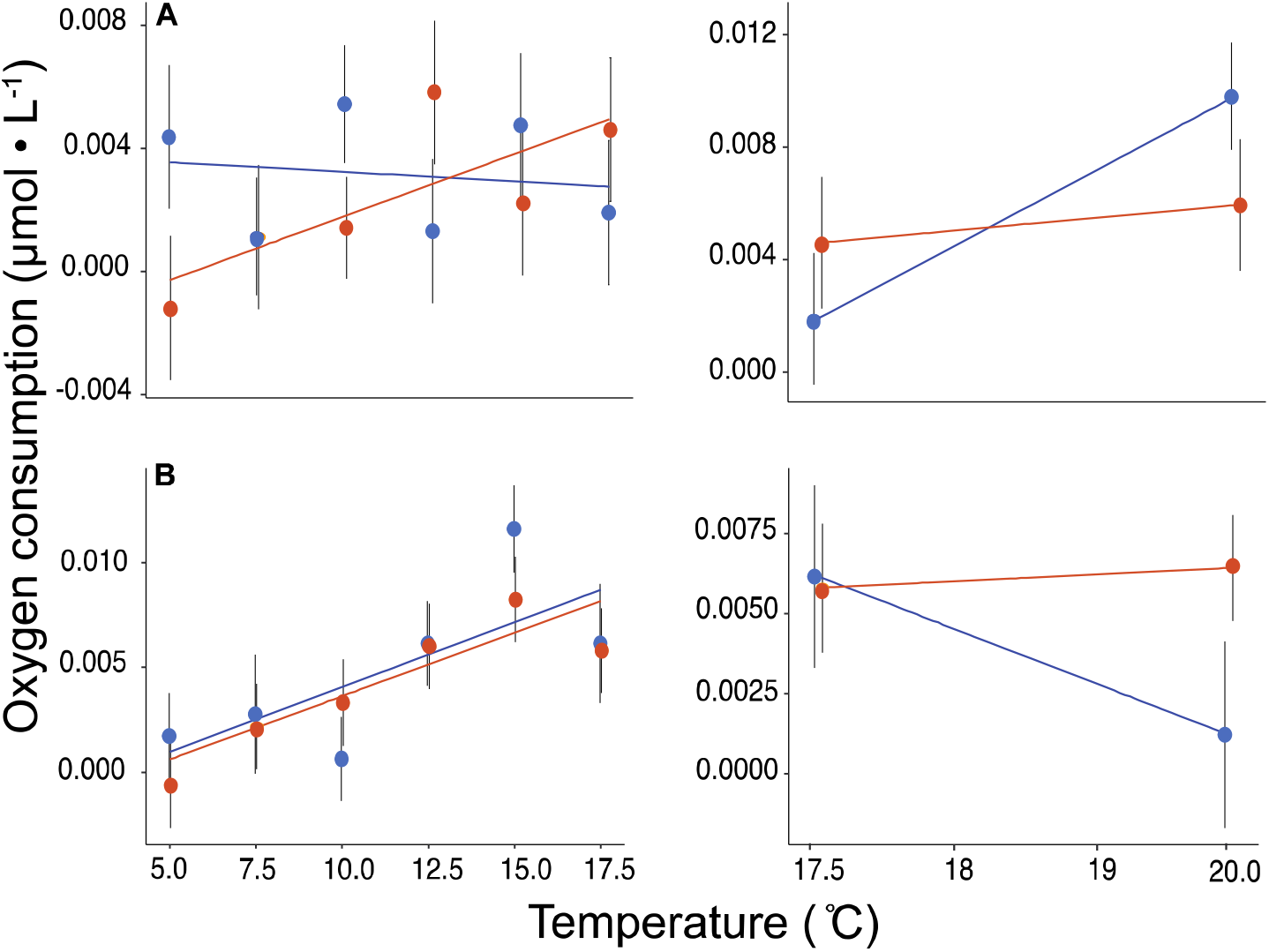
A comparison of the ascending (left panel) and descending (right panel) phases of metabolic profiles in perlid stoneflies at mid (A), and low (B) elevations. Plotted values are the least square means of metabolic rate in stoneflies from tropical (orange circles) and temperate (blue circles) streams, after accounting for body mass in the model. Vertical bars are standard errors.

When comparing descending phases, temperate mayflies had a marginally steeper decrease in metabolic rate at high elevation (Fig. 4A; *F*_(1,38)_ = 4.044, *p* = 0.051) but tropical mayflies had a much steeper decrease at low elevation (Fig. 4C; *F* _(1,28)_ = 9.464, *p* = 0.005). At mid elevation, there was no significant interaction (*F*_(1,35)_ = 2.477, *p* = 0.125; Fig. 4B). However, a Student’s t-test revealed that oxygen consumption values were significantly higher in mid-elevation tropical mayflies compared to temperate mayflies at 17.5 °C (*t* = 3.946, *p* = 0.001), but did not differ between latitudes at 20 °C (*t* = −1.006, *p* = 0.33). This suggests that the descending phase was steeper for tropical mayflies compared to temperate ones. Descending phases of stoneflies did not differ between tropical and temperate populations (Figs. 5 A & B).

Finally, maximum SMR was higher in tropical compared to temperate mayflies at all elevations (Fig. 6). Stoneflies showed an opposite pattern where tropical species had lower peaks at both measured elevations, although this was significant only at mid elevation (Fig. 6).

**Figure 6.**
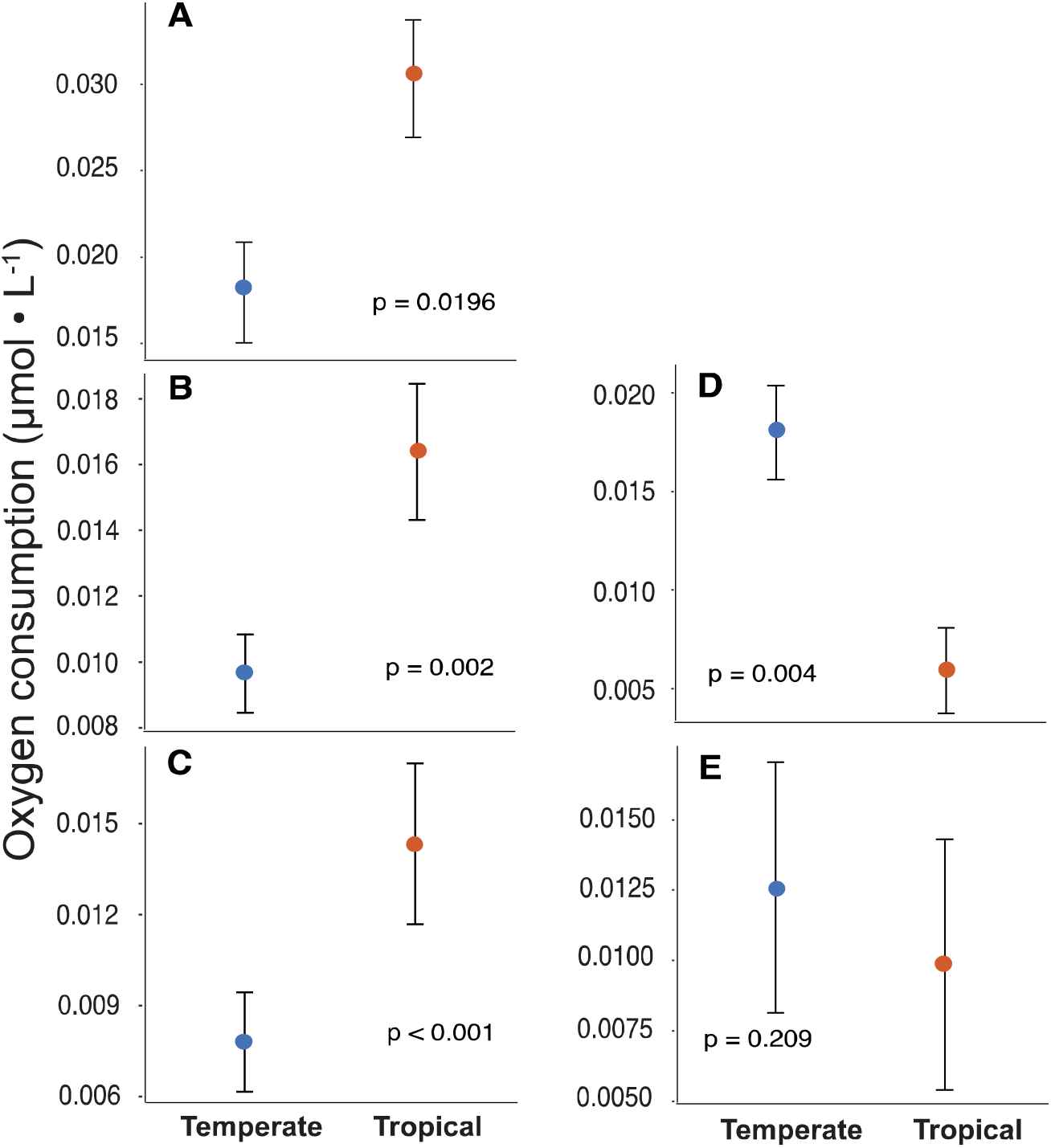
Peak SMRs (i.e., the highest metabolic rate) are compared between temperate and tropical baetid mayflies from high (A), mid (B), and low (C) elevations and perlid stoneflies from mid (D) and low (E) elevations. Plotted values are the least square means of metabolic rate for tropical (orange circles) and temperate (blue circles) insects, after accounting for body mass in the model. Vertical bars are standard errors and p values from an ANOVA for each elevation pair are shown.

### Thermal sensitivity based on Q_10_

When comparing across ‘stressful’ and ‘non-stressful’ temperatures, we found 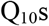 for both taxa to be higher at non-stressful temperatures than at stressful temperatures (Table 2). Across latitude, tropical mayflies showed generally greater sensitivity to temperature (higher 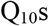) at both non-stressful and stressful temperatures than their temperate counterparts (Table 2; Fig. 7). Notably, at mid and low elevations, tropical mayflies were highly sensitive to temperature even at non-stressful temperatures (Q_10_ = 8.40 and 5.17, respectively). In contrast, tropical stoneflies were less sensitive than their temperate relatives for stressful and non-stressful temperatures (Fig. 7).

**Table 2.** Q_10_ values calculated for temperate and tropical mayfly and stonefly metabolic rates between typically non-stressful temperatures (7.5°C to 15°C) and typically stressful temperatures (15°C to 25°C).

|  |  | *Mayflies* |  | *Stoneflies* |  |
| --- | --- | --- | --- | --- | --- |
|  | Elevation | Temperate | Tropical | Temperate | Tropical |
| Non-stressful | High | 3.63 | 3.15 | n/a | n/a |
|  | Mid | 3.15 | 5.17 | 6.76 | 2.49 |
|  | Low | 2.36 | 8.40 | 6.66 | 5.78 |
| Stressful | High | 1.07 | 1.46 | n/a | n/a |
|  | Mid | 0.73 | 1.59 | 6.10 | 0.87 |
|  | Low | 1.34 | 1.67 | 1.25 | 0.42 |

**Figure 7.**
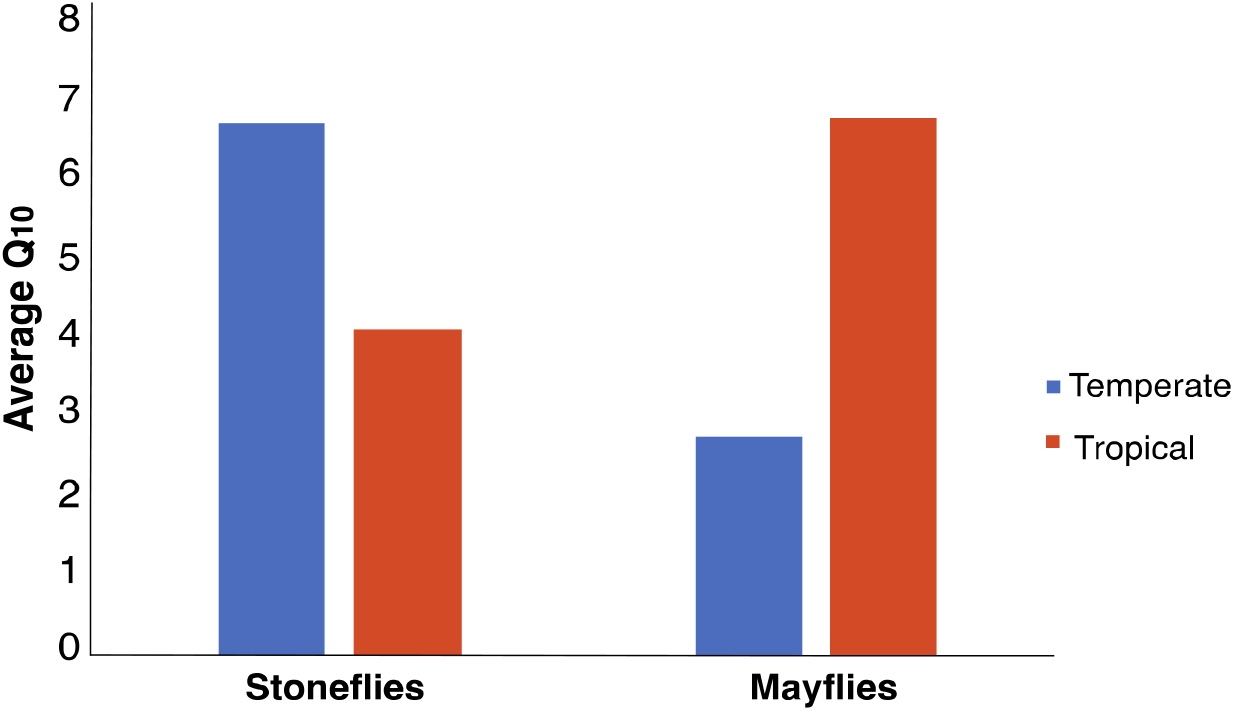
Comparison of average 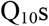 for mayflies and stoneflies during the ascending phase of the metabolic rate profiles.

## Discussion

Understanding the influence of temperature on the minimum energy demands of ectotherms is critical to predicting how global warming will impact different species (Huey & Kingsolver, 2019). Previous comparative studies of thermal tolerance (critical thermal limits) have generally found tropical ectotherms to be adapted to narrower ranges of temperatures compared to their temperate counterparts, suggesting that tropical species are more vulnerable to global warming (Deutsch et al., 2008; Gutiérrez-Pesquera et al., 2016; Shah et al., 2017b; Sunday, Bates, & Dulvy, 2011, 2012), a finding consistent with the CVH. Payne and Smith (2017) recently challenged such conclusions by arguing that narrower thermal breadth may simply reflect the acute effects of temperature on biological rates of organisms living in warmer environments. Resolving this debate requires an understanding of the acute effects of temperature on continuous biological rates like metabolism (Havird et al., 2020). Here, using a paired-elevation design, we compared the thermal sensitivity of standard metabolic rate (SMR) in closely related temperate and tropical aquatic insect nymphs. This study design enabled a comparison of closely-related temperate and tropical species under a common range of field, acclimation, and test temperatures. Our results did not support Payne and Smith’s (2017) model, but instead supported the CVH in mayflies, but not stoneflies.

Despite originating in streams with similar average environmental temperatures at the time of collection, tropical and temperate insects had different metabolic rate profiles (Figs. 2, 3). These results indicate that the variation between temperate and tropical species in our study does not simply reflect the acute effects of temperature on biological rates, but rather that there are physiological differences between taxa from these regions. Tropical baetid mayflies had metabolic rates that were more sensitive to temperature, and they exhibited stress at lower temperatures compared to their temperate counterparts. By contrast, tropical and temperate perlid stoneflies showed no strong divergence in their thermal sensitivity. Although the ecological implications of greater thermal sensitivity remain unknown for most ectotherms, in the absence of compensatory changes in productivity and energy expenditure, elevated metabolic rates could negatively impact the energy budget and capacity for aerobic performance in tropical mayflies by increasing baseline energetic costs (Dillon et al., 2010; Huey & Kingsolver, 2019).

### Comparison of metabolic rate profiles

In most cases, metabolic rate rose with increasing temperature, peaked, and then fell. This pattern, however, varied among taxa and populations and across elevations. As a group, the tropical mayflies are more thermally sensitive than their temperate counterparts (Figs. 2, 4, 6 and 7). The metabolic rate profiles of tropical mayflies had overall steeper ascending and descending phases, and higher 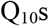. The particularly high 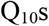 for mid and low elevation tropical mayflies were similar to those of North Atlantic stenothermal fishes when experiencing temperatures away from an optimum (Johnston, Clarke, & Ward, 1991). By contrast, high elevation tropical mayflies had similar 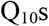 to their temperate relatives (Table 2), which may reflect the higher diel variation (i.e., variation across 24 h) experienced in streams above tree line in the Andes (higher standard error for stream temperature Table 1). In seasonally stable streams, such diel effects may play a potentially important role in regulating metabolic activity of aquatic insects (Gilchrist, 1995), but so far, no studies have explored the relative effects of diel and seasonal thermal variation in freshwater insects. In general, however, our results support the prediction that tropical mayflies, which experience more stable stream temperatures year-round (Shah et al., 2017b), are more sensitive to temperatures outside the ranges they normally experience.

Given that mayflies were paired by elevations having similar mean environmental and acclimation temperatures, these differences cannot be attributed to the acute or passive response of metabolic rate to higher temperatures alone (Havird et al., 2020). Instead, they appear to reflect divergence in thermal physiology as shown by the greater thermal sensitivity of metabolism in tropical mayflies. In contrast to tropical mayflies, temperate mayflies exhibited more modest increases in metabolism with temperature and had similar metabolic rate profiles across elevations (Fig. 2). We note that although there was no statistically significant interaction for the ascending limb between temperate and tropical mayflies at low elevation (Fig. 4), possibly driven by very low SMR at 7 ºC, tropical mayflies at low elevation exhibited strong stress responses to moderate temperatures (Fig. 2). Further, they have previously shown sensitivity to temperature with lower critical maximum temperatures and reduced acclimation responses (Shah et al., 2017a, 2017b). For both temperate and tropical mayflies, metabolic rates tended to be lower within the range of environmental temperatures they naturally experienced, suggesting there may be selection to reduce the minimum energetic costs associated with maintenance (Artacho & Nespolo, 2009; Bochdansky, Grønkjær, Herra, & Leggett, 2005; Brown, Marquet, & Taper, 1993). For example, even a moderate increase in stream temperature outside the normal range (Fig. 2, right edge of grey rectangles) is associated with peaks in metabolism, which has potentially cascading effects on energy budgets and possible chronic stress (Fig. 2). If selection acts to minimize SMR under typical stream temperatures (Brown et al., 1993), these results indicate that the narrower, more stable tropical stream temperatures make tropical mayflies more stenothermic and underlie their greater thermal sensitivity, as predicted by the CVH.

Such an interpretation is consistent with other aspects of temperate and tropical mayfly thermal physiology. First, thermal breadths (measured as the difference between critical thermal limits) reported in a parallel study were narrower for tropical (Ecuador) versus temperate (U.S.A) mayfly populations (Shah et al., 2017b). Second, in this study, we observed signs of stress in the metabolic rate profiles of tropical but not temperate mayflies. Specifically, after the decline in metabolic rate, secondary peaks occurred at 22.5 °C and 25 °C for most groups, which corresponded to observed signs of behavioral stress and mortality in tropical but not temperate mayflies. Secondary peaks are relatively rare in metabolic rate measurements but can occur, often near the critical thermal limits (Havird et al., 2020). Mayflies showed signs of behavioral stress at these high temperatures, including leg spasms, loss of righting, release from substrate, and mortality (see Supplemental Information Table S1, Figs. S3A, 3B). Collectively, these results support the idea that temperate mayflies are thermal generalists, with broad thermal breadths and reduced metabolic sensitivity to temperature, whereas tropical mayflies are thermal specialists, with narrower thermal breadths and increased metabolic sensitivity to temperature (c.f. Huey & Hertz, 1984).

The differences observed between temperate and tropical mayflies were mostly unambiguous and consistent with prior work in this system. Similarly, the *lack* of divergence between temperate and tropical stoneflies is also consistent with previous research on other physiological traits (Shah et al., 2017a, 2017b). Although there are differences in metabolic rate profiles between temperate and tropical stoneflies (e.g., higher metabolic rates at warmer temperatures in temperate but not tropical species), these differences are far less striking than those observed in mayflies. Ascending and descending phases did not differ between temperate and tropical stoneflies, but absolute maximum metabolic rates and 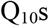 across non-stressful and stressful temperatures were higher in the temperate groups. Shah et al. (2017a) also found a lack of latitudinal differences in stoneflies when comparing acclimation abilities. One of the few other studies on the respiration rates of tropical stoneflies reported similar 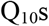 to those reported here for non-stressful temperatures in mid-elevation stoneflies (Jacobsen & Brodersen, 2008), corroborating our conclusions of lower thermal sensitivity in at least the *Anacroneuria* genus. Though many tropical species may be imperiled by warming (Deutsch et al., 2008; Dillon et al., 2010), the overall mixed support for this prediction in stoneflies cautions against generalizing about the vulnerability across tropical species.

Numerous hypotheses could explain the lack of divergence between temperate and tropical stoneflies. First, stoneflies as a group have a temperate origin and invaded tropical streams relatively recently (Baumann & Kondratieff, 1991; Stewart, Baumann, & Stark, 1974). Studies show that stoneflies are still adapting to tropical habitats (Polato et al., 2018), which may explain why thermal physiological traits in present-day temperate and tropical populations are similar.

Second, perlid stoneflies and baetid mayflies have different life histories, which may contribute to differential sensitivity to temperature. Stoneflies live much longer in streams as nymphs than mayflies (which have multiple generations per year), and may experience years of unpredictable variation in stream temperature (Clifford, 1982). In fact, many stoneflies enter diapause, unlike mayflies (Brittain, 1982; Hynes, 1976), giving them a greater ability to withstand adverse thermal conditions (Brittain, 1990). Once emerged, adult stoneflies often remain close to natal streams, whereas adult mayflies can disperse more widely in tropical (Finn, Encalada, & Hampel, 2016) and temperate (Polato et al., 2018) regions. Therefore, stoneflies rely more heavily on the nymph stage for dispersal (Brittain, 1990) and may experience more thermal variation while moving up- or downstream than juvenile mayflies. This increased likelihood of encountering thermal variation may favor the evolution of decreased thermal sensitivity (Muñoz & Bodensteiner, 2019).

Third, perlid stoneflies are important predators of baetid mayflies and midges (Peckarsky, 1980; Tamaris-Turizo, Turizo-Correa, & del Carmen Zúñiga, 2007) and they actively pursue prey in the wild (Peckarsky, 1980) and in laboratory settings (Sjöstrom, 1985; A.A.Shah *pers. obs*.). A reliance on aerobic metabolism and the need to maintain a high metabolic scope for active hunting (Kordas, Harley, & O’Connor, 2011) could constrain physiological divergence between temperate and tropical stoneflies. Unfortunately, few comparative studies of metabolic profiles are available to test the generality of such hypotheses. Nevertheless, higher maximum metabolic rates and higher 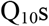 suggest some differences between temperate and tropical populations that could have ecological consequences when pursuing prey across a broad range of temperatures. Differences in thermal sensitivity between these predators and their prey may have community-level consequences under climate change (Gilman, Urban, Tewksbury, Gilchrist, & Holt, 2010; Grigaltchik, Ward, & Seebacher, 2012; Kordas et al., 2011; Pincebourde & Casas, 2019; Shah, Dillon, Hotaling, & Woods, 2020b; Vucic-Pestic, Ehnes, Rall, & Brose, 2011).

### Comparison of metabolic rates, critical thermal maxima, and performance curves as indicators of vulnerability

One should expect CT_MAX_, a commonly used metric for thermal sensitivity (Diamond et al., 2012; Lutterschmidt & Hutchison, 1997; Piyaphongkul, Pritchard, & Bale, 2012; Ribeiro, Camacho, & Navas, 2012), to coincide with the descending limb of the metabolic rate profiles (Fig. 1). Yet, previously measured values of CT_MAX_ in temperate mayflies occur at much higher temperatures (Fig.1A, B, C). For example, CT_MAX_ was ~29 °C in high elevation temperate baetid mayflies (Shah et al., 2017b) but they began to show elevated metabolic rates and behavioral signs of stress at lower temperatures (~17.5 °C). The large difference between CT_MAX_ and the onset of stress in this study is likely due to methodology, as rapid ramping or high starting temperatures can inflate estimated values of CT_MAX_ (Rezende, Tejedo, & Santos, 2011; Terblanche, Deere, Clusella-Trullas, Janion, & Chown, 2007). In addition, the discrepancy could also arise because CT_MAX_ and metabolic rates are governed by fundamentally different processes. CT_MAX_ is the result of changes in fluidity of cell membranes and whole-organism physiology in response to heat stress (Cossins, Friedlander, & Prosser, 1977; Hochachka & Somero, 2002), whereas standard metabolic rates arise from different activation energies of enzymes and sensitivity of biochemical processes to temperature (Rolfe & Brown, 1997). Thus, using CT_MAX_ in conjunction with other measures of thermal sensitivity such as metabolic rate provides a more complete assessment of thermal tolerance.

Another commonly measured indicator of thermal sensitivity is locomotor performance, in which T_OPT_ should lie at cooler temperatures than T-MR_PEAK_ (Fig. 1). However, in a separate study of swimming performance across temperature in the same mayfly species, we found no clear T_OPT_. Instead, TPCs for burst swimming were relatively flat in mayflies, with the exception of low elevation populations (Shah, Bacmeister, Rubalcaba, & Ghalambor, 2020a). Although it remains unclear why performance does not vary with temperature, one possible explanation is that selection favors a generalist strategy for swimming performance in all mayflies, perhaps because there is more thermal variation in streams than expected (Gilchrist, 1995). Alternatively, TPCs may not vary because burst swimming is less sensitive to temperature than more aerobically demanding activities like sustained swimming. Unlike most freshwater fishes, baetid mayflies do not engage in sustained swimming, but instead, walk and cling to rocks where their flattened bodies decrease shear stress from the water current (Weissenberger, Spatz, Emanns, & Schwoerbel, 1991). Thus, the relationship between temperature, SMR, and the allocation of energy to locomotion and other demands such as growth, reproduction, and maintenance remains an important area for future research.

### Climate change in Andean and Rocky Mountain Streams

Mountain stream temperatures are already increasing worldwide (Isaak & Rieman, 2013; Niedrist & Füreder, 2020). Temperatures of shallow mountain streams, such as those in this study, are also likely to show concomitant increases with air temperature (Birrell et al., 2020; Mohseni & Stefan, 1999; Pilgrim, Fang, & Stefan, 1998). Projected mean air temperature increases of 2-3 °C in the Andes and 3-4 °C in the northern Rocky Mountains by the 2050s (Nogués-Bravo, Araújo, Errea, & Martínez-Rica, 2007; Rangecroft, Suggitt, Anderson, & Harrison, 2016) are therefore troubling. However, previous analyses indicate that aquatic insects of the southern Rocky Mountains are relatively robust to warming because they may currently occur at suboptimal temperatures (Pyne & Poff, 2017). Moreover, mesocosm and field common garden experiments showed that temperate baetid mayflies grew faster when subjected to warmer water (A. Landeira-Dabarca et al.; A. Rugenski et al.; *unpubl. data*). By contrast, Andean mayfly growth declined with increasing temperature in mesocosms (A. Landeira-Dabarca et al.; *unpubl. data*) further supporting our conclusion that a warming environment will have greater negative impacts on tropical mayflies, especially at low elevation.

The effects of increasing temperatures may also lead to mismatches between dissolved oxygen supply and demand (Pörtner, 2001; Verberk, Bilton, Calosi, & Spicer, 2011; Verberk et al., 2016b). For many aquatic insects, the ability to maintain an adequate rate of oxygen uptake decreases at higher temperatures (Jacobsen & Brodersen, 2008; Rotvit & Jacobsen, 2013), and reduces energy available for activities like movement, growth, and egg development. Thus, even small increases in stream temperature can negatively impact insect metabolism and growth. This is illustrated in a field experiment, in which a moderate 2 °C temperature increase in a stream resulted in declines in abundance of Eurasian mayflies, presumably as a result of decreased oxygen in the stream (Verberk, Durance, Vaughan, & Ormerod, 2016a).

Finally, between the aquatic nymph stage and the flying adult stage there is potential for dispersal to mitigate the effects of future warming. Thermally sensitive tropical mayflies may seek cooler temperatures at higher elevations (Jacobsen, 2020), but whether the taxa in our study can move upstream as nymphs is debatable given the low dispersal rates of aquatic insects in these streams (Polato et al., 2018). Further, it is not known whether aquatic insects can mitigate the effects of warming by seeking out microhabitats that supply oxygen at high rates, i.e., those with relatively high oxygen concentrations or faster flows. Future work should therefore also focus on whether aquatic insects can move upstream to cooler waters or seek out high flow microhabitats to counteract the effects of warming in their native streams (Shah et al., 2020b; Sheldon, 2012).

## Acknowledgements

We thank the U.S. National Science Foundation for funding through the Dimensions of Biodiversity grant DEB-1046408; DEB-1045960, and DEB-1045991; and a Graduate Research Fellowship awarded to A.A.S under DGE-1321845. This research would not have been possible without the hard work of many field assistants. We especially thank M. Rojas, J. Fajardo, G. Waneka, B. Choat, L. Granizo, and L. Nagle for joining us over multiple field seasons. We are extremely grateful to J. Schreckinger, N. Quiroz, M. Celinscak, and E.J.G. Orejuela in Ecuador and K. L. Casner in the U.S. for help with field logistics. The Ministerio del Ambiente of Ecuador provided collection permits (Permit nos. 56-IC-FAU/FLO-DPN/MA and MAE-DNB-CM-2015-0017).

## Author contributions

A.A.S. and C.K.G. designed the experiments. A.A.S. conducted fieldwork and experiments, analyzed results, and wrote the manuscript with contributions from all authors.

## Data sharing and accessibility

Data generated in this study are openly available in the supporting information files of this article and on the public repository GitHub (https://github.com/Alisha-Shah/Standard-metabolic-rates.git).

